# Microbial metabolite *p*-cresol associates with constipation in adults with high-functioning autism spectrum disorder and induces constipation-like symptoms in mice

**DOI:** 10.64898/2025.12.31.697262

**Authors:** Narjis Kraimi, David Fabregat-Safont, Juliette Canaguier, Geoffroy Mallaret, Susana Barbosa, Julie Brouillet, Florian Lejuste, Alexandru Gaman, Jean-Romain Richard, Chema Sahnoun Derbel, Anouck Amestoy, Marion Leboyer, Oscar J. Pozo, Laetitia Davidovic

**Affiliations:** Institut de Pharmacologie Moléculaire et Cellulaire, Université Côte d’Azur, CNRS, INSERM, Valbonne, France; Applied Metabolomics Research Group, Hospital del Mar Research Institute, Barcelona, Spain; Environmental and Public Health Analytical Chemistry, Research Institute for Pesticides and Water (IUPA), Univ. Jaume I, Castelló, Spain; INSERM U955 IMRB, Translational Neuropsychiatry laboratory, AP-HP, DMU IMPACT, Fédération Hospitalo-Universitaire de Médecine de Précision en Psychiatrie (FHU ADAPT), Paris Est Créteil University, Créteil, France; Fondation FondaMental, Créteil, France; GHU Psychiatrie et Neurosciences Paris, Centre Médico-Psychologique de Belleville, Paris, France; Centre Hospitalier Charles-Perrens, Pôle Universitaire de Psychiatrie de L’enfant et de L’adolescent, CNRS, Aquitaine Institute for Cognitive and Integrative Neuroscience, INCIA, UMR 5287, Université de Bordeaux, Bordeaux, France

**Author notes:** Corresponding author: Laetitia Davidovic. Equal contribution (Narjis Kraimi, David Fabregat-Safont).

**Keywords:** ASD, microbiota, *p-*cresol, gastro-intestinal symptoms, cohort

## Abstract

Gastrointestinal (GI) symptoms are prevalent in Autism Spectrum Disorder (ASD), yet their biological correlates remain poorly understood. The gut microbial metabolite *p-*cresol is reported to be elevated in ASD, but its link to specific GI symptoms in adult high-functioning adults remains unexplored. We examined the relationship between serum levels of *p-*cresol and its conjugates (*p*-cresol sulfate, *p*-cresol glucuronide) with five common GI symptoms (diarrhea, constipation, abnormal stool aspect, bloating, abdominal pain) in 272 adults with high-functioning ASD. Clinical and dietary assessments were performed, and serum metabolites were measured using targeted metabolomics in 177 patients. Females with ASD had higher rates of GI symptoms and elevated *p-*cresol compared to males. Multivariable regression analysis showed that serum *p*-cresol, but not its conjugates, was selectively associated with constipation frequency, independent of sex, age, and diet. No associations were observed with the other GI symptoms. In mice, chronic *p*-cresol exposure induced constipation-like GI dysfunction, supporting a causal link. These findings identify *p*-cresol as a possible mediator of constipation in ASD. This work supports the development of tailored interventions for GI symptoms in adults with ASD.

**LAY SUMMARY:** People with autism spectrum disorder (ASD) often experience gastrointestinal (GI) symptoms, but the role of the gut microbiota remains unclear. Elevated levels of *p*-cresol, a gut microbiota–derived molecule, have been reported in ASD, though its link to specific GI symptoms, particularly in adults, remains unknown. We studied 272 adults with high-functioning ASD and we found that women with ASD reported more frequently GI symptoms and had higher levels of *p*-cresol than men. Higher *p*-cresol levels were associated with more frequent constipation, but not with other GI symptoms. In mice, *p*-cresol treatment induced signs of constipation, suggesting a causal role. Overall, our findings indicate that *p*-cresol may contribute to constipation in adults with ASD and support the need personalized medical approaches to digestive health in adults with ASD.

## INTRODUCTION

Autism Spectrum Disorder (ASD) is a complex neurodevelopmental condition characterized by early-onset difficulties in social communication, unusually restricted, repetitive behaviors, and atypical sensory processing (Lai et al., 2014). Gastrointestinal (GI) symptoms such as constipation, diarrhea, abdominal pain, and bloating are frequent comorbidities in ASD and their severity correlates with the severity of behavioral symptoms (Croen et al., 2015; Gorrindo et al., 2012; Holingue et al., 2018; Lasheras et al., 2023; Leader et al., 2021; Mazurek et al., 2013; McElhanon et al., 2014; Nikolov et al., 2009). This high prevalence of GI symptoms has intensified interest in the gut-brain axis and its potential role in ASD pathophysiology (Vuong & Hsiao, 2017). Alterations in gut microbiota composition and microbial metabolite production may contribute to both GI and behavioral symptoms in ASD (Needham et al., 2020; Vuong & Hsiao, 2017). Also, ASD-related restricted interests drive severe food selectivity and a less-diverse diet, further reducing microbial diversity and impacting GI motility (Yap et al., 2021).

Among microbial metabolites, *p*-cresol (pC) has gained particular attention due to its reported elevation in urine and feces of individuals with ASD and possible association with ASD and GI symptom severity, in particular constipation (Altieri et al., 2011; De Angelis et al., 2013; Flynn et al., 2025; Gabriele et al., 2014; Gevi et al., 2016; Kang et al., 2018; Persico & Napolioni, 2013; Piras et al., 2022; Victoria-Montesinos et al., 2025). However, large multivariable analyses rigorously adjusting for confounders such as sex, age, or diet are generally lacking to document the links between pC levels and ASD. In preclinical work, we have shown that chronic pC exposure in mice alters gut microbiota composition and induces ASD-like behaviors, including social behavior deficits and stereotypies (Bermudez-Martin et al., 2021; Canaguier et al., 2025; Mallaret et al., 2025), but pC impact on GI function has not yet been explored. pC is mainly produced in the colon from dietary tyrosine by gut bacteria (Challand et al., 2010; Saito et al., 2018), absorbed, and subsequently conjugated to *p*-cresol sulfate (pCS) and *p-*cresol glucuronide (pCG) in the liver, with contribution from kidney and intestinal mucosa (Blachier & Andriamihaja, 2022). These conjugated forms, together with remaining free pC, circulate systemically and are predominantly excreted renally (Gryp et al., 2017). Despite extensive work on urinary pC and its conjugates in ASD, circulating blood levels of pC and its conjugates in ASD remain poorly characterized, although they may more directly reflect systemic exposure and detoxification capacity.

These research gaps to link GI symptoms to their biological correlates in ASD are compounded by broader biases. Adults remain underrepresented despite ASD being a lifelong condition with both core associated symptoms persisting across lifespan. Only a few studies have examined GI symptoms in adults with ASD (Croen et al., 2015; Lasheras et al., 2023; Leader et al., 2021; Torenvliet et al., 2025) and their biological underpinnings are still poorly understood in this population. Moreover, most biomarker studies in ASD have included children with intellectual disability, although epidemiological data indicate that approximately 50-60% of individuals with ASD have average to above-average intelligence and are therefore high-functioning (Polyak et al., 2015). This bias has significant implications, as high-functioning individuals may present with different patterns of comorbidities compared to those with intellectual disability (Howes et al., 2018). In addition, factors known to influence microbial composition, metabolite production, and host conjugation, such as sex, age, and diet, are not consistently accounted for in clinical studies, even though sex differences in ASD presentation and dietary selectivity are well documented (Wenzell et al., 2024; Werling & Geschwind, 2013).

To address these gaps, the present study quantifies serum pC and its conjugates (pCS, pCG) in adults with high-functioning ASD and examines associations between these metabolites and GI symptoms controlling for age, sex, and diet. In parallel, we investigate the impact of chronic pC exposure on GI function in mice, thereby integrating human and preclinical data to elucidate potential causal links between microbial pC, GI dysfunction, and ASD.

## METHODS

### HUMAN STUDY

#### Study sample

A total of 272 patients were enrolled under informed consent. Main inclusion criteria included being above 18-year-old, diagnosed with ASD Diagnostic and Statistical Manual of Mental Disorders IVth edition (DSM-IV) and having an intellectual quotient (IQ) above 70 based on the Weschler Adult Intelligence Scale (WAIS-II and IV). ASD core symptoms were assessed by trained psychiatrists with clinical expertise using the Autism Diagnostic Observation Scheduled (ADOS), the Autism Diagnostic Interview Revised (ADI-R), and the DSM-IV classification. Data on socioeconomic status and race/ethnicity were not recorded. The study sample characteristics are presented in Table 1. A subsample of 177 patients agreed to provide serum samples. Veinous blood was sampled in collection tubes, allowed to coagulate for 30 min. at RT, and centrifuged within 2 hrs (1,500g, 15 min., 4°C). Serum was stored at - 80°C until use.

**Table 1.**
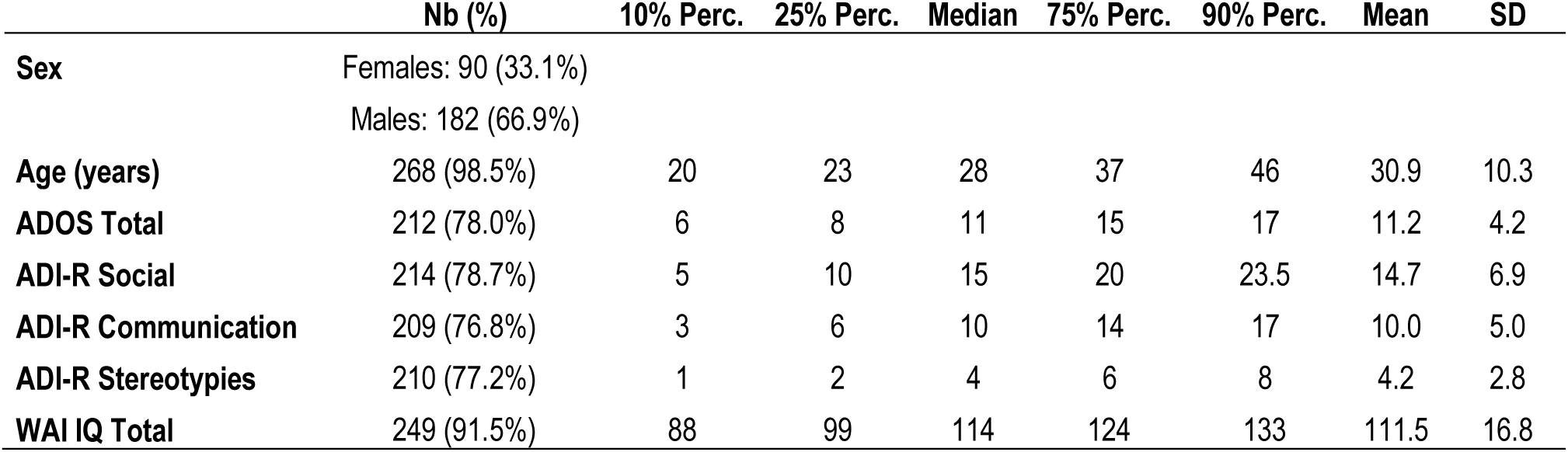
Study sample.

|  | Nb (%) | 10% Perc. | 25% Perc. | Median | 75% Perc. | 90% Perc. | Mean | SD |
| --- | --- | --- | --- | --- | --- | --- | --- | --- |
| **Sex** | Females: 90 (33.1%)<br>Males: 182 (66.9%) |  |  |  |  |  |  |  |
| **Age (years)** | 268 (98.5%) | 20 | 23 | 28 | 37 | 46 | 30.9 | 10.3 |
| **ADOS Total** | 212 (78.0%) | 6 | 8 | 11 | 15 | 17 | 11.2 | 4.2 |
| **ADI-R Social** | 214 (78.7%) | 5 | 10 | 15 | 20 | 23.5 | 14.7 | 6.9 |
| **ADI-R Communication** | 209 (76.8%) | 3 | 6 | 10 | 14 | 17 | 10.0 | 5.0 |
| **ADI-R Stereotypies** | 210 (77.2%) | 1 | 2 | 4 | 6 | 8 | 4.2 | 2.8 |
| **WAI IQ Total** | 249 (91.5%) | 88 | 99 | 114 | 124 | 133 | 111.5 | 16.8 |

### GI symptoms

GI symptoms in adulthood were assessed using a structured questionnaire specifically designed for adults with ASD, rather than standard tools such as the Rome IV criteria or the GSRS (Drossman & Hasler, 2016; Svedlund et al., 1988). Rome IV emphasizes categorical diagnoses and may miss subthreshold GI symptoms common in ASD. The GSRS places substantial emphasis on upper-GI complaints that are less central to our research question. Our 5-item instrument focuses on the core lower-GI symptoms frequently observed in ASD (diarrhea, constipation, abnormal stool characteristics, bloating and abdominal pain), uses simple, concrete wording, and is administered by a trained physician ensuring comprehension. This format reduces cognitive and attentional burden, improves reliability in individuals with communication difficulties, and yields quantitative, symptom-based scores that are well suited for association analyses. Patients were asked how often, during adulthood, they had experienced episodes (lasting at least two consecutive weeks) of: 1. Abnormally loose or watery stools; 2. Constipation, defined as fewer than two bowel movements per week or abnormally hard stools; 3. Abnormally foul-smelling stools or stools of unusual colour (e.g., with pus or blood); 4. Significant abdominal bloating; 5. Significant abdominal discomfort or pain. Each item was rated on a 5-point Likert scale (0 = never, 1 = rarely, 2 = sometimes, 3 = often, 4 = very often). Although the GI symptom outcomes were collected on a 5-point Likert scale and are therefore strictly ordinal, we first modelled them as a numerical variable under the assumption that adjacent categories reflect approximately equal intervals along an underlying continuous latent trait, a common and empirically supported practice in epidemiological and psychometric analyses that yields minimally biased estimates while improving statistical efficiency and interpretability of regression coefficients (Norman, 2010; Sullivan & Artino, 2014).

### Diet

Diet intake was assessed with a short food frequency questionnaire based on the French NutriNet-Santé cohort (European Clinical Trials Database 2013-000929-31). 154 participants (56.61% of the original sample) agreed to report their usual frequency of consumption of 29 predefined food groups over the previous year on a 6-point Likert-type scale (0 = never or almost never; 1 = <1 time/week; 2 = about once/week; 3 = 2–3 times/week; 4 = 4–6 times/week; 5 = once/day or more), which we modelled as numerical variables. The full list of food items is provided in Supplementary Table S1. To reduce dimensionality and capture overall dietary patterns, we performed a principal component analysis (PCA) on the consumption frequencies of the 29 items (after centering and scaling). PCA on the 29 food items yielded four components (eigenvalues > 1) explaining 41.95% of the total variance (Supplementary Table S1). Inspection of the loading structure showed that food groups with high loadings were heterogeneous, with recommended and non-recommended items loading together on the same components. Therefore, no coherent, easily interpretable dietary patterns could be identified. Component scores for PC1 and PC2 were nevertheless used as empirical summary variables to adjust for overall dietary variation. Full PCA results, including eigenvalues, loadings, and variable contributions, are presented in Supplementary Table S1.

### Determination of *p*-cresol, *p-*cresol sulfate, and *p-*cresol glucuronide in serum

Serum free pC was quantified using a targeted liquid chromatography-tandem mass spectrometry (LC-MS/MS) targeted method after derivatization with 1,2-dimethylimidazole-5-sulfonyl chloride (5-DMISC), which provides superior analytical performance compared with dansyl chloride and 1,2-dimethylimidazole-4-sulfonyl chloride (Fabregat-Safont et al., 2025). The method achieved a limit of detection of 4 pg/mL and a limit of quantification of 20 pg/mL, enabling measurement of low-abundance free pC in human serum. Details of sample preparation and instrumental conditions have been reported previously (Fabregat-Safont et al., 2025).

A second targeted LC-MS/MS method was developed to quantify the two pC conjugates (pCS, pCG), in the same samples with limits of quantification of 1 ng/mL. Chromatographic conditions matched those used for free pC, but a dilute-and-shoot sample preparation was applied. Briefly, 20 µL of serum were mixed with 80 µL of a methanolic internal standard solution containing isotopically labelled p-cresol sulfate-d₄ and p-cresol glucuronide-d₇, vortexed and centrifuged. The supernatant was injected (3 µL) into the LC-MS/MS system. Calibration curves were prepared in methanol by mixing 20 µL of calibrator with 80 µL of the internal standard mixture. Limits of quantification were 1 ng/mL for both conjugated metabolites. Two selected-reaction monitoring transitions were acquired for each analyte: pCS (negative ion mode, m/z 187→80 and 187→107) and pCG (positive ion mode, m/z 302→194 and 302→141).

### Statistical analyses

Comparison analyses of variables were performed using the Wilcoxon-Mann & Whitney U test for numerical variables or with the Chi-square test for categorical variables. Correlations between numerical variables and assimilated were performed using Spearman’s ρ correlation coefficient rank test with Benjamini & Hochberg’s multiple testing correction for each block of analysis. Statistical significance was set at an adjusted p-value < 0.05.

To meet normality assumptions, pCG levels were log-transformed. To address skewness and stabilize variance, variables related to pC levels, pCS levels, constipation frequency, and age were subjected to Box-Cox transformation (Y = (Y^ λ - 1) / λ (Box & Cox, 1964)). The optimal λ Box-Cox parameter for the transformation was determined using maximum likelihood estimation to ensure approximate normality of the variables and homoscedasticity of the residuals to result an optimal fit against the normal distribution (pC: λ = 0.35; pCS: λ = 0.43; Age: λ = −0.76; 1+constipation score: λ = −0.59) (Supplementary Table S2).

Multivariable linear or logistic regressions were then conducted to study the determinants of pC, pCG and pCS levels and their associations with constipation (either considering the full range of the Lickert scale or dichotomizing frequent/very frequent (scores 3, 4) vs. the rest (scores 0, 1, 2) as outcomes). Age, sex, and dietary patterns were included as covariates in the models. Model diagnostics were performed to confirm that the assumptions of linearity, normality of residuals, and absence of multicollinearity were satisfied. The regression coefficients and 95% confidence intervals (CI) are reported on the transformed scale since both constipation and pC were Box-Cox transformed. Significance was set at p<0.05. We did not adjust for multiple testing since regressions were considered independently.

All statistical analyses were performed using GraphPad Prism 10.4.1 and R,.

## ANIMAL STUDY

### Ethics statement for animal housing and experimentation

Animal housing and experimentation were conducted in facilities certified by regional authorities. The study was conducted in accordance with procedures approved by the local ethics committee for animal experimentation (Ciepal-Azur) and the Ministère de l’Enseignement Supérieur et de la Recherche (APAFIS #21355-2019062414391395 v3, #51032-2024082612133352 v5).

### Animals and *p*-cresol treatment

Weaned C57BL/6J male mice (21-28 day-old) were obtained from Charles River (France). Upon arrival, animals were randomly allocated to experimental groups and housed in open cages of 4 to 5 animals in a temperature (22–24 °C) and hygrometry (70–80%)-controlled room with a 12 h light/dark cycle (lights on from 7:00 a.m. to 7:00 p.m.) and *ad libitum* access to water and food (standard chow).

pC (Sigma-Aldrich) was dispensed in sterile drinking water at a concentration of 0.25 g/L for 4 weeks. Based on a mean body mass of 25 g and a mean drinking water consumption of 5 mL/24 h, this is equivalent to a dose of 50 mg/Kg/24 h *per os*. It has previously been reported that this treatment was not interfering with basic physiological parameters (drink/food consumption, body weight) but induced social interaction deficits and stereotypies, with no impact on locomotor activity, exploratory behavior, anxiety or cognition (Bermudez-Martin et al., 2021; Mallaret et al., 2025).

### Fecal output and fecal dry matter

Mice were placed individually empty cages for 1 hour, with *ad libitum* access to food and water. Feces were collected immediately after defecation using forceps and placed into pre-weighed individual tubes. Fecal pellet number and weight were recorded prior lyophilization (VirTis® AdVantage lyophilizer) to determine fecal dry matter.

### GI transit time

Total intestinal transit time was assessed using carmine red, which is not absorbed by the intestinal epithelium. Animals were not fasted prior to the experiment and had access do drinking water and food. Mice received by gavage a solution of 0.5% methylcellulose (w/v) containing 3.75% (w/v) Carmine red (final dose 0.3 mg/g body weight) or not (vehicle control group). Gavage time was recorded as T0. Feces were monitored regularly for the appearance of red-colored pellets. Total GI transit time was defined as the interval between T0 and the first observation of carmine red-containing feces.

### Colonic motility

Colonic motility was evaluated *in vivo* using the glass bead expulsion test. Mice were lightly anesthetized with isoflurane, and a glass bead (3 mm diameter; Fisher Scientific) was gently inserted 2 cm into the distal colon using a lubricated metallic catheter. The animals were then placed individually in clean cages without bedding, with free access to food and water. The colonic transit time was defined as the time interval between bead insertion and bead expulsion, with a cut-off at 60 min.

### Intestinal permeability

Fluorescein-5-(and-6)-sulfonic acid (FSA, Invitrogen) and horseradish peroxidase (HRP, Sigma Aldrich) were used as paracellular and transcellular permeability marker, respectively. Solutions of FSA and HRP were prepared at concentration of. Mice received 2 mg/g of body weight of each compound dissolved in saline solution (10 mg/mL each). After 1.5 hours, 100 µL of blood was collected from the tail tip into EDTA-coated tubes. Plasma was isolated by centrifugation (5000 g, 5 min) at 4°C, and stored at −80°C.

For FSA quantification, plasma was diluted 1:50 in PBS and transferred to a 96-well plate. Fluorescence intensity was measured using a microplate reader (BioTek Cytation 5) with excitation at 488 nm and emission at 518 nm. For HRP measurement, 40 µL of plasma diluted 1:30 in PBS was mixed with 100 µL of 3,3’,5,5’-tetramethylbenzidine (TMB). Once the solution turned blue, reaction was stopped by adding 100 µL of 1N H_2_SO_4_. Absorbance was then read at 450 nm.

### Determination of circulating *p*-cresol

Mice were euthanized by a lethal injection of pentobarbital and blood sampled by cardiac puncture, transferred to heparin-coated tubes (BD Microtainer PST Tubes Lithium Heparin). The tubes were gently inverted 5 times and after 30 min at RT, plasma was collected by centrifugation (10,000g, 2 min, RT), snapped-frozen in liquid nitrogen, and stored at – 80 °C until analysis with the method described above for human samples (Fabregat-Safont et al., 2025).

### GI tract segments histological preparations

The abdominal cavity was opened and the portion of GI tract excised by cutting between the rectum and the stomach pylorus. GI tissue was thoroughly unfolded and wrapped around a Whatman cardboard piece fixation for 24 h with 4% paraformaldehyde at 4°C. GI tissue was rinsed 3 times with PBS, unfolded and the small intestine divided into duodenum, jejunum, and ileum, while the colon was kept intact. Each segment was cut into 1-cm pieces, stacked, and wrapped tightly in microporous tape to form bundles as described (Williams et al., 2016). Bundles were embedded in paraffin. Sections (5 µm) were cut on a microtome, mounted on Superfrost Plus slides, and incubated for 1 h at 65°C. Section were deparaffinized, subjected to hematoxylin and eosin staining, and imaged using a Leica DMD108 digital microscope. For each mouse, we quantified manually the number of the different cell types in at least 3 villi and 3 crypts, across three different images per GI segment (duodenum, jejunum, ileum, colon), as recommended in (Williams et al., 2016). The ImageJ software was used to measure villi and crypts lengths.

### Statistical analyses

Regarding animal data analyses, if n<10, comparisons between two independent groups were performed using the two-tailed Mann–Whitney U test. If n>10, data were first tested for normality using the Kolmogorov-Smirnov test. When the assumption of normality was met, comparisons between two independent groups were performed using the two-tailed Student t test. When normality was not met in either one of the groups or both, comparisons between two independent groups were performed using the two-tailed Mann–Whitney U test. All statistical analyses were performed using GraphPad Prism 10.4.1, with significance set at p<0.05.

## RESULTS

### Study sample

Our study sample consisted in N=272 adults (30.9 ± 10.3 years) diagnosed with ASD (Table 1), 33.1% (N=90) of which were females. These patients presented an average ADOS score of 11.2 (± 4.2). The average ADI-R subscores related to social reciprocity, communication, and stereotypies/restricted interests were respectively 14.7 (± 6.9), 10 (± 5), and 4.2 (± 2.8) which corresponds to the range of ADI-R scores found in the literature for ASD patents (Pereira et al., 2018; Ren et al., 2023). Patients did not exhibit comorbid intellectual disability, with IQ above 70 and a mean IQ of 111.5 (± 16.8). A subsample of 251 patients out of 272 (93.3% of the original study sample) reported on their GI symptoms. Bloating and abdominal pain were the most frequently reported GI symptoms among adult ASD patients, followed by constipation, diarrhea, and abnormal stool aspect (Table 2). More specifically, patients reported very frequent episodes of bloating (14.4%), abdominal pain (8%), constipation (7.6%), diarrhea (7.6%), or abnormal stool (1.2%). Female ASD patients reported more frequently episodes of diarrhea, constipation, abdominal bloating, and abdominal pain than male patients, while abnormal stool aspect was equally reported (Table 2).

**Table 2.**
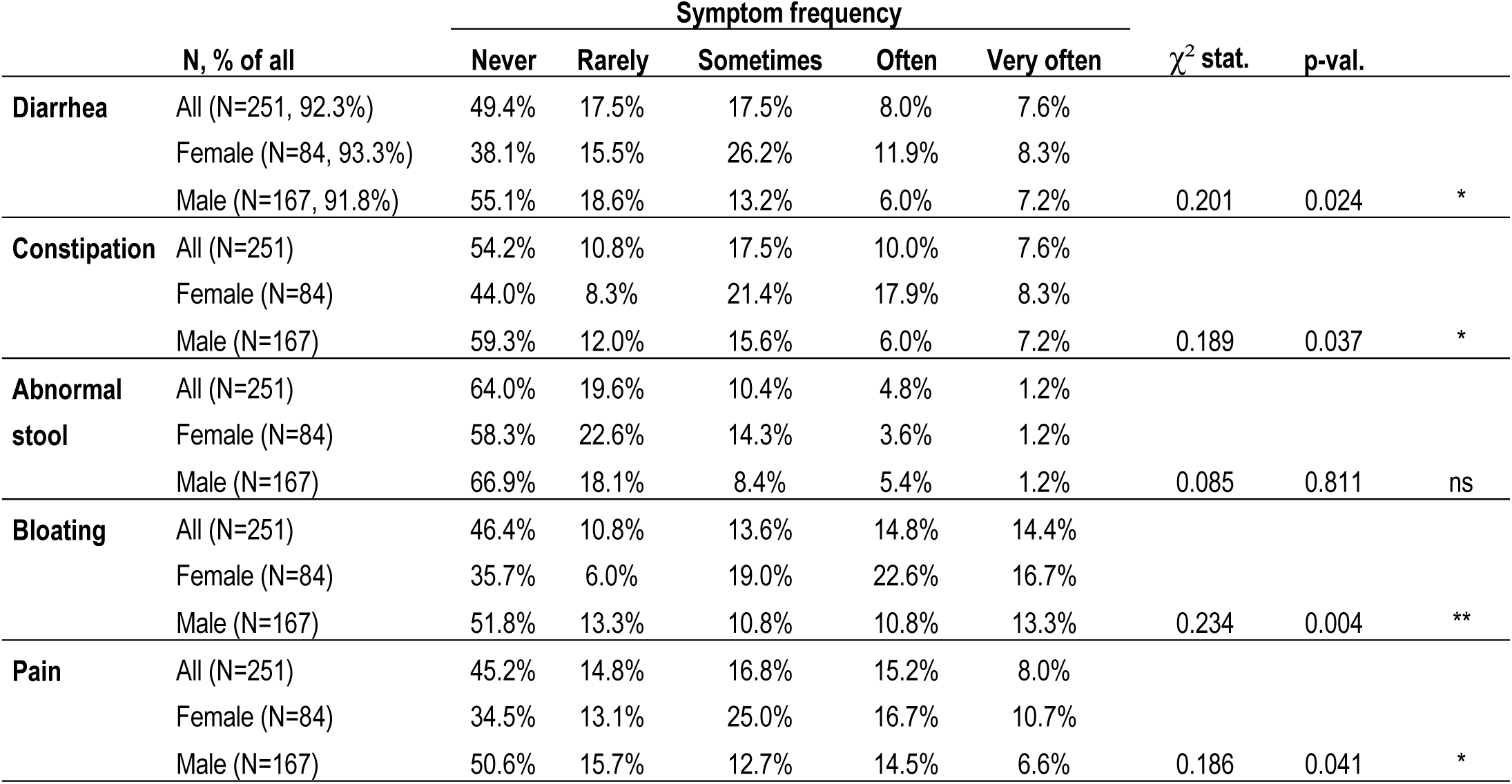
GI symptomatology in ASD patients and stratified by sex. Chi-square-test to test the effect of sex: *, p<0.05; **, p<0.01; ns, p>0.05.

### Circulating levels of *p*-cresol and its conjugates *p*-cresol glucuronide and *p*-cresol sulfate and influence of sex, age, and diet

We measured the levels of pC and its main conjugates pCG and pCS in a subsample of 177 patients who had agreed for serum sample collection (65% of the original study sample). pC was detected in the serum in the pg/ml range, while pCG and pCS were much more abundant in the ng/ml range (Figure 1, Table 3). Free unconjugated pC represented only 0.009% (± 0.02%) of total serum pC, while pCG represented 0.5% (± 0.05%) and pCS 99.5% (± 0.5%) (Supplementary Table S3). Finally, pCG accounted for less than 0.51% (± 0.0055%) of *p-*cresol sulfate (Supplementary Table S3). Spearman’s correlation analysis indicated medium correlations between pC levels and pCG (ρ=0.389, p<0.0001) or pCS (ρ=0.366, p<0.0001) levels and strong correlations between the levels of its conjugates pCG and pCS (ρ=0.878, p<0.0001) (Figure 1D, Supplementary Table S4).

**Figure 1.**
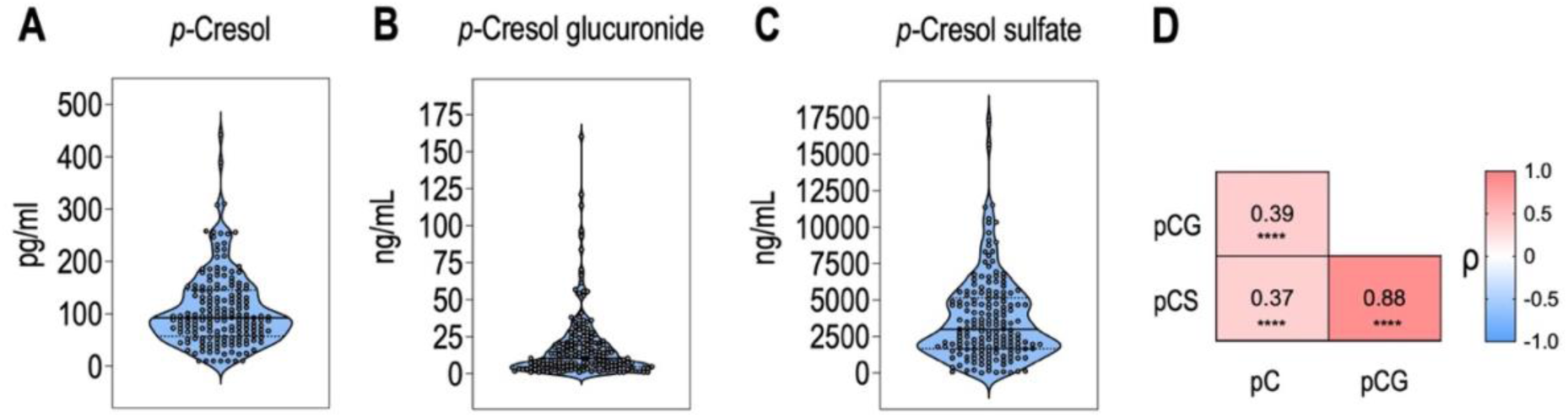
Circulating *p*-cresol, *p-*cresol glucuronide, and *p*-cresol sulfate levels in ASD patients. **A.** Serum *p*-cresol levels. **B.** Serum *p*-cresol glucuronide levels. **C.** Serum *p*-cresol sulfate levels. **D.** Spearman’s correlations between serum levels of *p*-cresol (pC), *p*-cresol glucuronide (pCG), *p*-cresol sulfate (pCS). ****, p<0.0001. **A-C** Data are presented as violin plots showing distribution quartiles and all values as dots (N=177).

**Table 3.** Serum *p*-cresol (pC), *p*-cresol glucuronide (pCG), and *p*-cresol sulfate (pCS) levels in ASD patients and stratified by sex. Mann-Whitney U test to test the effect of sex: **, p<0.01; ns, p>0.05.

|  |  | LLOD | Min. | 10% Perc. | 25% Perc. | Median | 75% Perc. | 90% Perc. | Max. | Mean | SD | p- value | Significance |
| --- | --- | --- | --- | --- | --- | --- | --- | --- | --- | --- | --- | --- | --- |
| pC (pg/ml) | All (N=177) |  | 10 | 34.64 | 57.1 | 92.8 | 145.3 | 188.8 | 442 | 107.2 | 70.5 |  |  |
|  | Female (N=62) | 10 | 30.8 | 46.84 | 71.8 | 107.2 | 172.6 | 242.4 | 442 | 129.5 | 80.4 |  |  |
|  | Male (N=114) |  | 10 | 25.8 | 51.25 | 87.4 | 128.8 | 176.9 | 310.2 | 94.88 | 61.34 | 0.003 | ** |
| pCG (ng/ml) | All (N=176) |  | 1 | 2.066 | 4.695 | 10.62 | 21.25 | 36.63 | 160 | 17.86 | 22.75 |  |  |
|  | Female (N=62) | 1 | 1 | 2.509 | 5.903 | 14.71 | 24.85 | 55.23 | 160 | 21.8 | 28.54 |  |  |
|  | Male (N=114) |  | 1 | 1.65 | 3.758 | 8.77 | 20.67 | 36.22 | 113.4 | 15.57 | 18.72 | 0.072 | ns |
| pCS (ng/ml) | All (N=176) |  | 17 | 757.2 | 1672 | 2999 | 5151 | 7006 | 17301 | 3670 | 2865 |  |  |
|  | Female (N=62) | 10 | 52 | 1043 | 1744 | 3288 | 4911 | 7350 | 15669 | 3829 | 2844 |  |  |
|  | Male (N=114) |  | 17 | 641 | 1561 | 2778 | 5281 | 7124 | 17301 | 3568 | 2892 | 0.428 | ns |

We then studied the factors that could influence pC, pCG, and pCS levels, including age, sex, and diet. When stratifying by sex, pC levels were on average 36% higher in females than in males, while pCG and pCS levels were similar in female and male subjects (Table 3). No correlations were observed between pC, pCG, or pCS levels and age or dietary patterns captured by the first 4 PC of the food frequency questionnaire PCA analysis (Supplementary Table S5).

We then utilized multivariable linear regression to study altogether the influence of diet, age and sex on pC levels. This showed a significant influence of sex, with pC levels more elevated in female subjects, while there was no effect of age or diet on pC levels (Table 4). As for pCG or pCS levels, multivariable analyses confirmed that there were no significant effects of sex, age, or diet (Supplementary Tables S6, S7).

**Table 4.** Effects of sex, age and diet on serum *p*-cresol levels. *p-*Cresol levels and age were Box-Cox transformed prior to analysis; estimates and confidence intervals (CI) are reported on the transformed scale. **, p<0.01; ns, p>0.05.

| Variable | Estimate | 95% CI | p-value | Significance |
| --- | --- | --- | --- | --- |
| Sex[Female] | 2.036 | [0.689 ; 3.383] | 0.0034 | ** |
| Age | -1.220 | [-31.32 ; 28.89] | 0.9361 | ns |
| Diet-PC1 | -0.032 | [-0.247 ; 0.184] | 0.7712 | ns |
| Diet-PC2 | 0.108 | [-0.128 ; 0.345] | 0.3656 | ns |
| Diet-PC3 | 0.206 | [-0.05 ; 0.462] | 0.1141 | ns |
| Diet-PC4 | -0.018 | [-0.282 ; 0.247] | 0.894 | ns |

### Associations between serum *p-*cresol, *p-*cresol glucuronide and *p-*cresol sulfate with GI symptoms in high-functioning individuals with ASD

We then studied the relationships between pC, pCG and pCS levels and the severity of GI symptoms. No correlations were observed between pC, pCG or pCS with the frequency of diarrhea, abnormal stool aspect, bloating, or abdominal pain (Supplementary Table S8). Though, we highlighted a significant positive correlation between pC levels and increased frequency of constipation episodes, which survived adjustment for multiple testing (ρ=0.237, adj. p=0.018). This correlation was only observed for free pC and not for its conjugates pCG or pCS (Supplementary Table S8).

Multivariable linear regression showed that pC levels were positively associated with increased frequency of constipation episodes, adjusting for sex and age (model a, Table 5, Supplementary Table S9). In a sensitivity analysis, we also tested the associations further adjusting for diet, using diet PC1 and PC2 and found that pC levels were significantly associated with more frequent constipation episodes (model b, Table 5, Supplementary Table S9). In a third analysis, we dichotomized constipation frequency score between frequent/very frequent vs the never/sometimes/often group. The association between constipation and pC levels were also retained in this model (model c, Table 5, Supplementary Table S9). These effects were specific to pC as no associations were observed with its conjugates pCS or pCG (Supplementary Tables S10, S11).

**Table 5.** Associations between p-cresol and constipation frequency. *p-*cresol levels, age, and constipation frequency scores were Box-Cox transformed prior to analysis; estimates and confidence intervals (CIs) are reported on the transformed scale. Back-transformed estimates for *p*-cresol are provided in Supplementary Table S9. Model a: Linear regression with constipation frequency as a continuous outcome (5-point Likert scale), adjusted for sex and age. Model b: Linear regression with constipation frequency as a continuous outcome (5-point Likert scale), adjusted for sex, age, and dietary patterns (PC1, PC2). Model c: Logistic regression with constipation frequency dichotomized (frequent/very frequent [scores 3–4] vs. never/sometimes/often [scores 0–2]), adjusted for sex, age, and dietary patterns (PC1, PC2). *, p<0.05; **, p<0.01; ns, p>0.05.

|  | Variable | Estimate | 95% CI | p-value | Significance |
| --- | --- | --- | --- | --- | --- |
| **Model a** | **pC** | **0.066** | **[0.019 ; 0.112]** | **0.006** | **\*\*** |
|  | Sex [Female] | 0.190 | [0.120 ; 0.501] | 0.227 | ns |
|  | Age | 1.444 | [-5.275 ; 8.162] | 0.672 | ns |
| **Model b** | **pC** | **0.080** | **[0.026 ; 0.134]** | **0.004** | **\*\*** |
|  | Sex [Female] | 0.260 | [-0.121 ; 0.640] | 0.170 | ns |
|  | Age | -5.803 | [-13.94 ; 2.338] | 0.160 | ns |
|  | Diet-PC1 | 0.005 | [-0.054 ; 0.064] | 0.869 | ns |
|  | Diet-PC2 | -0.034 | [-0.099 ; 0.0304] | 0.296 | ns |
| **Model c** | **pC** | **0.189** | **[0.006 ; 0.387]** | **0.043** | **\*** |
|  | Sex [Female] | 1.019 | [-0.035 ; 2.318] | 0.060 | ns |
|  | Age | -17.38 | [-46.93 ; 10.42] | 0.222 | ns |
|  | Diet-PC1 | -0.120 | [-0.348 ; 0.091] | 0.269 | ns |
|  | Diet-PC2 | -0.040 | [-0.284 ; 0.193] | 0.739 | ns |

### *p*-cresol exposure induces constipation-like phenotypes in mice

Since the observed association between circulating pC and constipation does not, on its own, establish causality, we directly tested this in mice which we treated with pC dissolved in drinking water. A 4-week pC exposure significantly increased circulating pC levels by approximately 2-fold compared to controls (Supplementary Figure S1), in agreement with our anterior work. Histological assessments indicated similar intestinal epithelial architecture across groups, showing that chronic pC exposure unlikely compromises gut epithelium integrity (Supplementary Figure S2). Comprehensive *in vivo* GI phenotyping revealed a constipation-like phenotype in pC-treated mice. Relative to controls, mice chronically exposed to pC exhibited reduced hourly fecal output and increased fecal dry matter content (Figure 2 A, B). Colonic motility did not differ (Figure 2 C) and whole gut transit time was unchanged (Figure 2 D). In contrast, both paracellular and transcellular permeability to tracer molecules significantly increased pC-treated mice (Figure 2 E, F).

**Figure 2.**
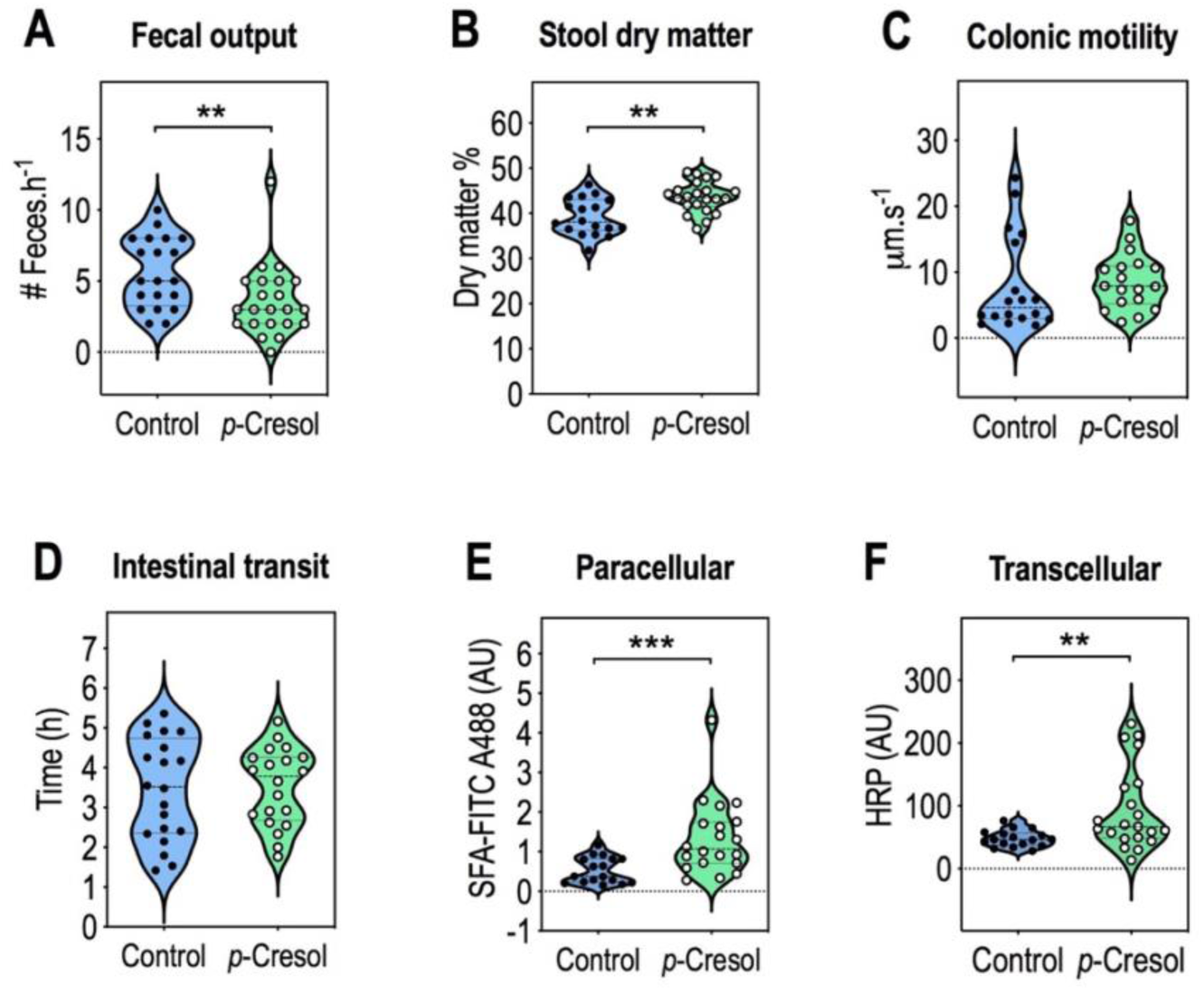
Mice exposed to *p*-cresol exhibit constipation-like phenotypes and increased intestinal permeability. **A.** Fecal output with the number of feces expelled per hour. Two-tailed Mann–Whitney U test, **p=0.007. **B.** Stool dry matter (%). Unpaired Student-t test, ** p=0.001. **C.** Colonic motility (µm. s^-1^). Two-tailed Mann–Whitney U test, p=0.16. **D.** Intestinal transit time (h). Unpaired Student-t test, p=0.865. **E.** Paracellular permeability (SFA-FITC A488 (AU)). Two-tailed Mann-Whitney U test, ***p=0.0005. **F.** Transcellular permeability (HRP (AU)). Unpaired Student-t test, ** p=0.007. Data are presented as violin plots showing distribution quartiles and individual values plotted as dots (n=17-20 Control, n=18-22 *p*-Cresol).

## DISCUSSION

Our study clarifies the circulating levels of the microbial metabolite *p*-cresol and its conjugates in adults with ASD and provides convergent human and mechanistic evidence that links *p*-cresol with constipation. We report three principal findings: i) circulating unconjugated *p*-cresol is present at very low abundance compared with its conjugates, with *p*-cresol sulfate overwhelmingly predominant; ii) free p-cresol, but not *p*-cresol sulfate or *p-*cresol glucuronide, associates with the frequency of constipation episodes, independently of age, sex, and diet, and iii) chronic GI exposure to *p*-cresol in mice induces a constipation-like phenotype, supporting a causal relationship.

### GI symptomatology and circulating levels of *p*-cresol and its conjugated metabolites

In adults with high-functioning ASD, we found that episodes of constipation, diarrhea, bloating, and abdominal pain were more frequently reported in female individuals than in males, in agreement with a recent study showing that females were more prone to bowel conditions (Torenvliet et al., 2025). Regarding more specifically constipation, 7.6% of individuals with ASD experienced very frequently constipation. This is consistent with a previous study reporting that 4.45% of adults with an ASD diagnosis (based on the International Classification of Diseases) had constipation syndrome, compared to only 1.39% in sex- and -aged matched healthy individuals without such diagnosis (Croen et al., 2015). Regarding free pC, it was present in low abundance (pg/ml range) while pCG was present in the ng/ml range, the dominant circulating form was pCS, in agreement with previous studies showing that pC is predominantly sulfated to pCS, with only a minor glucuronidated fraction (Gryp et al., 2017; Serrano-Tomás et al., 2025). Only two studies have previously quantified the circulating levels of pCS and pCG in the plasma of ASD patients (Kang et al., 2020; Needham et al., 2021), both in pediatric cohorts and using relative quantification, precluding direct comparison with our data. We found that free pC correlated modestly with its conjugates, whereas both conjugates were strongly correlated. Conjugation processes by sulfotransferases and UDP-glucuronyltransferases are efficient and occur rapidly, likely limiting the free pC in the bloodstream. Notably, ASD has been associated with global impairment in sulfation capacity, including reduced phenol sulfotransferase activity, compared with unaffected siblings or neurotypical individuals (Pagan et al., 2021). Such deficit could decrease pC sulfo-conjugation and increase pC bioavailability. These findings underscore the importance of quantitative methods that distinguish free from conjugated metabolites when studying microbial metabolites in ASD.

### Sex, but not age or diet, as modulating factor of *p-*cresol levels in adult ASD individuals

We observed a significant effect of sex on circulating free pC, with higher levels in females ASD patients. This observation aligns with prior studies showing sex-based differences in gut microbiota composition and metabolic activity, which may be influenced by hormonal factors such as estrogen (Hokanson et al., 2024). We found no effect of age on pC levels in adult individuals with ASD, suggesting relative stability in adulthood. At first glance this may appear inconsistent with findings in a cohort of ASD individuals aged 2-18 years where elevated pC was primarily in small children (age 2-7) (Altieri et al., 2011). However, this previous study measured total pC (free plus conjugates) and did not use multivariable statistical analyses to isolate the effect of age, limiting comparability. These considerations raise the possibility that gut dysbiosis and its metabolic consequences can be chronic features of ASD, irrespective of age. Contrary to expectations, dietary patterns were not significantly associated with levels of pC or its conjugates, whether considering each of the 29 diet items individually or the first four diet PC. This differs from studies linking extreme dietary exposures (e.g. protein supplements, vegetarian diet) either to increased or decreased pC sulfate production in healthy volunteers (Geypens et al., 1997; Patel et al., 2012). However, these are extreme dietary habits which may not be reflected in our study sample. These results highlight the complexity of diet-microbiome-metabolites interactions and argue for more detailed dietary assessments in future work.

### Association between free *p*-cresol levels and constipation and causal support from adult mice

We show that higher frequencies of constipation episodes were specifically associated with higher circulating free pC levels and this association persisted after adjustment for sex, age, and diet. No association was detected for circulating pCS and pCG. This selectivity could be explained by the fact that prolonged colonic stasis during constipation episodes can enhance luminal pC production (Al Hinai et al., 2019). Importantly, plasma pC sulfate levels were reduced in individuals who have undergone colectomy, as compared to individuals with a colon (Aronov et al., 2011), suggesting that the colon contributes substantially to the generation of its precursor pC. One interpretation could be that constipation promotes luminal pC biosynthesis, and, consequently, systemic accumulation of pC in ASD. Free pC could also potentially alter epithelial water exchanges and stool consistency, as it was previously shown to decrease transepithelial electrical resistance and increase paracellular permeability in human intestinal cell monolayers (Wong et al., 2016). In line with this, our mouse experiments provide complementary causal evidence of a constipation-like phenotype induced by chronic pC exposure in adult mice. First, decreased fecal pellet count and harder feces are two well-known hallmark features in validated murine models of constipation (Ge et al., 2017; Liang et al., 2016; Lin et al., 2021). The absence of an effect on whole-gut transit time and colonic motility suggests a functional dysregulation of colonic fluid handling and epithelial barrier integrity rather than a global hypomotility phenotype. A lack of gut transit delay has been also reported at early stages of constipation, when stool hardening and increased intestinal permeability can occur without reduced motility (Liang et al., 2016). Consistently, we found that pC increased both paracellular and transcellular intestinal permeability without overt mucosal injury across the duodenum, jejunum, ileum, and colon, indicating subtle biochemical or molecular disruption rather than overt structural damage. This aligns with prior *in vitro* work showing that pC disrupts gut epithelial junctions and increases permeability (Andriamihaja et al., 2015; Blachier & Andriamihaja, 2022; Flynn et al., 2025; Wong et al., 2016; Zhang et al., 2025). Moreover, patients with constipation exhibit increased intestinal permeability as suggested by increased serum ovalbumin concentrations (Khalif et al., 2005), and mice colonized with feces from patients with constipation displayed abnormal defecation parameters and reduced mucin expression levels, suggesting gut barrier alteration (Cao et al., 2017). Our experimental data in mice support the notion that pC could causally contribute to constipation in ASD.

### Study limitation

GI symptoms were assessed using a physician-administered questionnaire rather than standardized Rome IV criteria, this may have introduced misclassification and limits comparability with other studies. pC, pCS, and pCG levels were assessed in a subsample of 177 patients for which serum samples were available, future longitudinal on larger samples would strengthen the findings. Preclinical experiments were performed in male mice only, whereas in our human cohort females showed higher pC levels and GI symptom burden. This sex mismatch limits direct translational inference and highlights the need to include female animals in future mechanistic work.

## CONCLUSIONS

Free pC, unlike its conjugated forms, associates with constipation frequency in adults with high-functioning ASD, suggesting that GI-focused assays should prioritize measuring the unconjugated compound. Furthermore, if free pC contributes to constipation, interventions reducing its luminal production or enhancing elimination warrant evaluation in ASD, including interventions with dietary fibers or pre/probiotics previously shown to lower pC (De Preter et al., 2007; Nakabayashi et al., 2011; Salmean et al., 2015). Finally, sex should be incorporated into clinical assessment and trial design, as females showed higher free pC and more frequent GI symptoms. Although based on retrospective symptoms and a single cohort, these findings highlight free pC as a clinically relevant microbial metabolite and support testing microbiota-targeted treatments. These results provide a foundation for future research into gut-microbiota brain interactions in ASD and their therapeutic implications.

## Acknowledgements

We thank the IPMC Animal Facility for expert animal care and advice, in particular Dr. Thomas Lorivel and Pauline Pozzo di Borgo.

## Author Contributions

None.

## STATEMENTS AND DECLARATIONS

### Ethical considerations

This clinical study was approved by French regulatory authorities (Comité de Protection des Persones Ile de France II) on Nov. 20^th^, 2017 (#RCB/EUDRACT: 2017-A01318-45). CODECOH approval to use biological samples was received 2021, May 27^th^ (#DC-2021-4457). All participants provided informed consent for data and biological samples’ collection.

### Consent to participate

Not applicable.

### Consent for publication

Not applicable.

### Declaration of conflicting interest

The author(s) declared no potential conflicts of interest with respect to the research, authorship, and/or publication of this article.

### Funding statement

NK was funded by a postdoctoral fellowship from the Fondation pour la Recherche Médicale (ARF202409019389). DFS acknowledges his postdoctoral grant (CD24/00027), funded by Instituto de Salud Carlos III and co-funded by the European Union (Fondo Social Europeo Plus (FSE+)). JC was funded by a PhD fellowship from the Fondation pour la Recherche Médicale (ECO202006011591). JB was funded by a PhD fellowship from the EUR LIVE Graduate School of Research (UPEC). We acknowledge the financial contribution of the Agence Nationale de la Recherche (ANR-19-CE14-0024-MicrobiAutism to ML and LD, ANR-2024-CE18-1909 SynBact4Autism to LD).

### Data availability

The datasets supporting the conclusions of this article are included within the article (and its additional files) and other data will be made available upon reasonable request to LD or ML via the FondaMental Asperger Expert Centers. Notably, the clinical dataset is available in Supplementary Table 2.

## Notes

### Competing Interest Statement

The authors have declared no competing interest.

