## Supplementary data for "Microbial metabolite *p*-cresol associates with constipation in adults with high-functioning autism spectrum disorder and induces constipation-like symptoms in mice"

**Supplementary tables:** **10**

**Supplementary figures:** **2**

**SUPPLEMENTARY TABLES**

**Table S1. Principal component analysis of dietary components.**

*Click here to access/download supplementary xls file*

**Table S2. Complete dataset and transformed data.**

*Click here to access/download supplementary xls file*

**Table S3. Serum *p*-cresol, *p*-cresol glucuronide, and *p*-cresol sulfate levels, and relative ratios in ASD patients.**

*Click here to access/download supplementary xls file*

**Table S4. Correlations between serum *p*-cresol, *p*-cresol glucuronide, and *p*-cresol sulfate levels in ASD patients.** Spearman's ρ correlation coefficient rank test with Benjamini & Hochberg's multiple testing correction.

| | Spearman's ρ | | | p-val | | | adj. p-val | | |
|---|---|---|---|---|---|---|---|---|---|
| | pC | pCG | pCS | pC | pCG | pCS | pC | pCG | pCS |
| **pC pg/ml** | 1 | 0.389 | 0.366 | NA | 9.11E-08 | 5.27E-07 | NA | 2.73E-07 | 1.58E-06 |
| **pCG ng/ml** | 0.389 | 1 | 0.878 | 9.11E-08 | NA | 5.48E-58 | 2.73E-07 | NA | 1.65E-57 |
| **pCS ng/ml** | 0.366 | 0.878 | 1 | 5.27E-07 | 5.48E-58 | NA | 1.58E-06 | 1.65E-57 | NA |

**Table S5. Correlations between age and diet-derived principal components and serum *p*-cresol, *p*-cresol glucuronide, and *p*-cresol sulfate levels in ASD patients.** Spearman's ρ correlation coefficient rank test with Benjamini & Hochberg's multiple testing correction.

| | Spearman's ρ | | | p-val | | | adj. p-val | | |
|---|---|---|---|---|---|---|---|---|---|
| | pC | pCG | pCS | pC | pCG | pCS | pC | pCG | pCS |
| Age | 0.002 | 0.044 | 0.046 | 0.974 | 0.565 | 0.543 | 0.974 | 0.974 | 0.974 |

| | | | | | | | | | |
|---|---|---|---|---|---|---|---|---|---|
| Diet-PC1 | 0.024 | -0.009 | 0.043 | 0.807 | 0.926 | 0.657 | 0.974 | 0.974 | 0.974 |
| Diet-PC2 | 0.027 | -0.073 | -0.129 | 0.782 | 0.452 | 0.178 | 0.974 | 0.974 | 0.974 |
| Diet-PC3 | 0.093 | -0.028 | -0.033 | 0.336 | 0.775 | 0.731 | 0.974 | 0.974 | 0.974 |
| Diet-PC4 | -0.008 | -0.112 | -0.013 | 0.930 | 0.244 | 0.893 | 0.974 | 0.974 | 0.974 |

**Table S6. Effects of sex, age and diet on serum *p*-cresol glucuronide levels.** ns, p>0.05.

| Variable | Estimate | 95% CI | p-value | Significance |
|---|---|---|---|---|
| Sex[Female] | 0.1426 | [-0.05209 ; 0.3372] | 0.1494 | ns |
| Age | 0.7774 | [-3.572 ; 5.127] | 0.7237 | ns |
| Diet-PC1 | -0.007469 | [-0.03863 ; 0.02369] | 0.6355 | ns |
| Diet-PC2 | -0.01294 | [-0.04709 ; 0.02121] | 0.4541 | ns |
| Diet-PC3 | 0.01456 | [-0.02245 ; 0.05157] | 0.437 | ns |
| Diet-PC4 | -0.02422 | [-0.06245 ; 0.01400] | 0.2117 | ns |

**Table S7. Effects of sex, age and diet on serum *p*-cresol sulfate levels.** ns, p>0.05.

| Variable | Estimate | 95% CI | p-value | Significance |
|---|---|---|---|---|
| Sex[Female] | 2.243 | [-8.211 ; 12.70] | 0.6713 | ns |
| Age | 43.34 | [-190.3 ; 277.0] | 0.7136 | ns |
| Diet-PC1 | 0.09224 | [-1.581 ; 1.766] | 0.9132 | ns |
| Diet-PC2 | -1.018 | [-2.853 ; 0.8156] | 0.2733 | ns |
| Diet-PC3 | 0.2249 | [-1.763 ; 2.213] | 0.8229 | ns |
| Diet-PC4 | -0.6328 | [-2.686 ; 1.420] | 0.5423 | ns |

**Table S8. Correlations between serum *p*-cresol, *p*-cresol glucuronide, and *p*-cresol sulfate levels and the severity of GI symptoms.** Spearman's ρ correlation coefficient rank test with Benjamini & Hochberg's multiple testing correction.

| | ρ | | | p-val | | | adj. p-val | | |
|---|---|---|---|---|---|---|---|---|---|
| | **pC** | **pCG** | **pCS** | **pC** | **pCG** | **pCS** | **pC** | **pCG** | **pCS** |
| **Diarrhea** | -0.048 | -0.183 | -0.182 | 0.552 | 0.022 | 0.023 | 0.845 | 0.088 | 0.088 |
| **Constipation** | 0.237 | 0.016 | 0.046 | 0.003 | 0.846 | 0.570 | 0.018 | 0.932 | 0.845 |
| **Abnormal stool** | 0.031 | -0.001 | 0.004 | 0.704 | 0.989 | 0.963 | 0.845 | 0.889 | 0.963 |
| **Bloating** | 0.068 | 0.051 | 0.075 | 0.399 | 0.530 | 0.354 | 0.598 | 0.699 | 0.607 |
| **Abdominal pain** | -0.110 | -0.051 | -0.059 | 0.175 | 0.525 | 0.469 | 0.350 | 0.699 | 0.607 |

**Table S9. Back-transformed log estimates and 95% CI for associations between *p*-cresol levels with constipation frequency.** **p<0.01; *p<0.05

| | Variable | Estimate | 95% CI | p-value | Significance |
|---|---|---|---|---|---|
| **Model a** | **pC** | **1.024** | **[1.007 ; 1.043]** | **0.006** | ** |
| **Model b** | **pC** | **1.032** | **[1.010 ; 1.052]** | **0.004** | ** |
| **Model c** | **pC** | **1.070** | **[1.002 ; 1.150]** | **0.043** | * |

**Table S10. Associations between *p*-cresol sulfate with constipation frequency.** Model adjusted for sex, age. ns, p>0.05.

| Variable | Estimate | 95% CI | p-value | Significance |
|---|---|---|---|---|
| pCS | 0.002076 | [-0.00345 ; 0.0076] | 0.4593 | ns |
| Sex [Female] | 0.2895 | [-0.01996 ; 0.5989] | 0.0665 | ns |
| Age | 1.060 | [-5.836 ; 7.955] | 0.7619 | ns |

**Table S11. Associations between *p*-cresol glucuronide with constipation frequency.** Model adjusted for sex, age. ns, p>0.05.

| Variable | Estimate | 95% CI | p-value | Significance |
|---|---|---|---|---|
| pCG | 0.04865 | [-0.2558 ; 0.3531] | 0.7527 | ns |
| Sex [Female] | 0.2857 | [-0.02553 ; 0.5970] | 0.0717 | ns |
| Age | 1.729 | [-5.739 ; 8.075] | 0.7387 | ns |

### SUPPLEMENTARY FIGURES

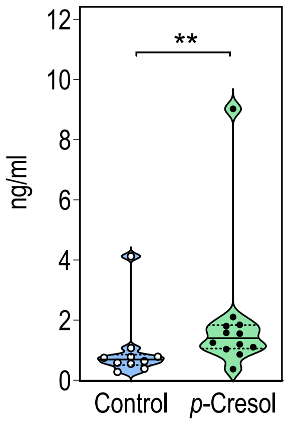


**Supplementary Figure S1. Mice exposed to *p*-cresol exhibit increased circulating levels of *p*-cresol**

Plasma levels of *p*-Cresol. Two-tailed Mann–Whitney U test. **p = 0.009; n=10 Control, n=12 *p*-Cresol.

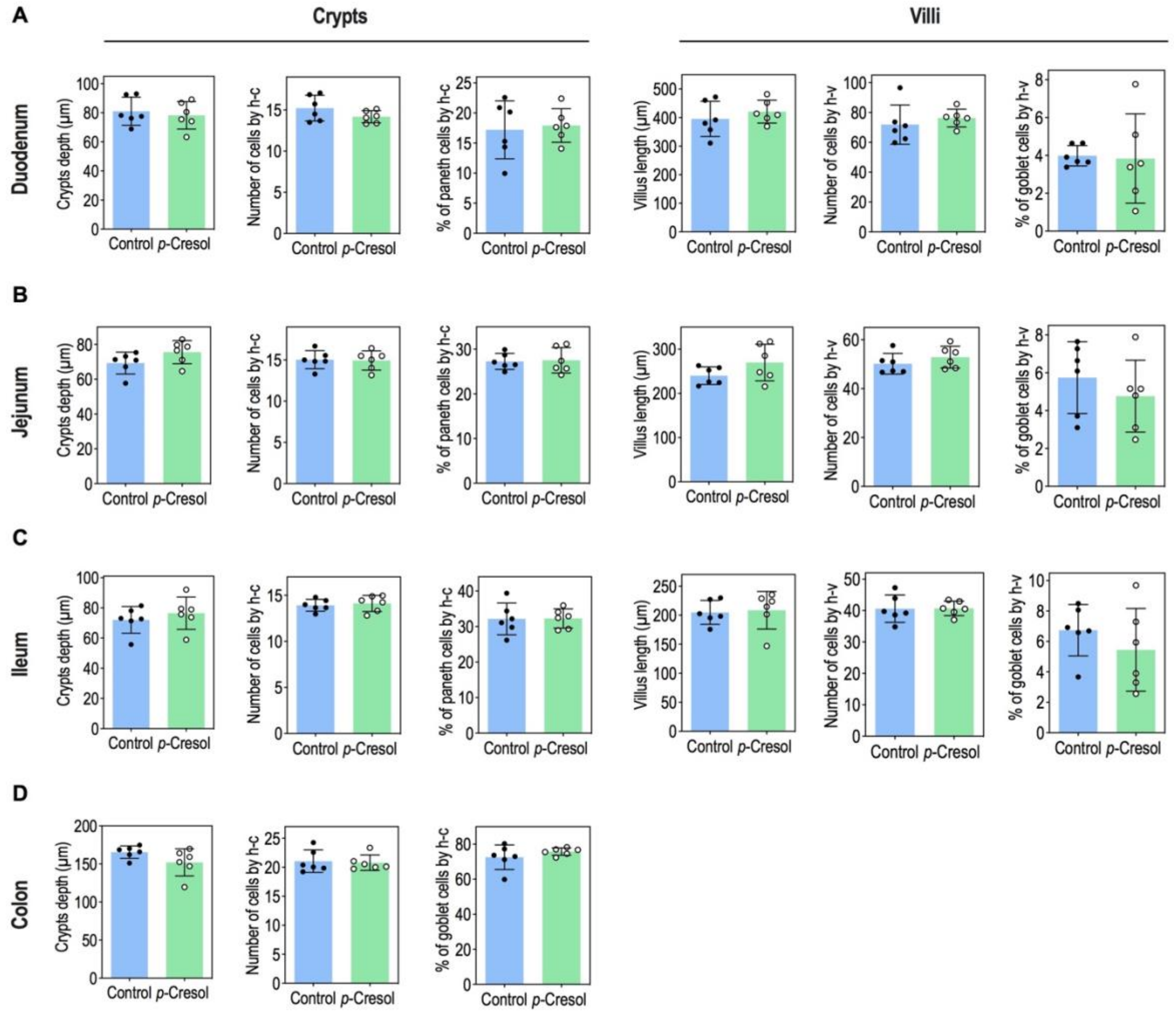


**Supplementary Figure S2. Histological measurements in intestinal tissues from control and *p*-cresol–treated mice.**

**A.** Measures of crypts depth (µm), total number of cells in crypts, % of Paneth cells in crypts, villus length (µm), total number of cells in villi, and % of goblet cells in villi in duodenum. **B.** Measures of crypts depth (µm), total number of cells in crypts, % of Paneth cells in crypts, villus length (µm), total number of cells in villi and % of goblet cells in villi in jejunum. **C.** Measures of crypts depth (µm), total number of cells in crypts, % of Paneth cells in crypts, villus length (µm), total number of cells in villi, and % of goblet cells in villi in ileum. **D.** Measures of crypts depth (µm), total number of cells in crypts and % of goblet cells in crypts in colon section. **A-D**. Data are presented as dot plots showing means ± standard deviation (n=6 Control. n=6 *p*-Cresol, for each animal, three images/GI segment and three villi or crypts/image analyzed). Two-tailed Mann–Whitney U test. $p > 0.05$.
